# Climate Change Impacts on *Broussenetia papyrifera* Pollens - Metabolome Investigations and Prospects of Allergy Prevalence

**DOI:** 10.1101/2022.02.14.480393

**Authors:** Muhammad Humayun, Saadia Naseem, Richard E. Goodman, Zahid Ali

**Author notes:** **Corresponding author:**, Plant Biotechnology and Molecular Pharming Lab, Department of Biosciences, COMSATS, University Islamabad (CUI), Pakistan.

## Abstract

*Broussonetia papyrifera* (*B. papyrifera*) is an allergenic plant in the mulberry family that grows at varied elevations and climatic conditions worldwide. In northern Pakistan, *B. papyrifera* is abundant and it produces a substantial amount of pollen that disperses in the air causing allergies in some humans. Climate change affects pollen production. To investigate potential changes in pollens development and potential allergenicity, *B. papyrifera* pollens were collected in summer and in spring from different regions in Pakistan. Study samples were subjected to morphological analysis, Fourier Transform Infrared Spectroscopy (FTIR) analysis for biochemical differences, and Liquid Chromatography-Mass Spectrometry (LCMS) for metabolome analysis. Morphological studies of the dried pollen by light microscopy showed seasonal and regional differences in pollens size and exine morphology. FTIR analysis showed inter-regional and inter-seasonal differences in the metabolome of the pollen. Differences in lipid and protein functional groups of pollen from different regions showed variation in the FTIR spectra. These differences in FTIR spectra correlated with the changing climatic conditions. Metabolome analysis of targeted pollen samples identified 33 organic compounds of seven different groups. Four unsaturated fatty acids were identified that have a potential role in allergic responses. The findings in this study are unique in demonstrating climatic variables that effect *B. papyrifera* pollen physiology (FTIR analysis) which also confirms differences in pollen-associated lipid metabolites identified by LCMS analysis. These results demonstrate information that may be used to predict potential changes in allergy risks from pollens of *B. papyrifera* in the future. The findings may provide a model for predicting variation in pollen structure and associated allergies in response to climate changes for other species.

## Introduction

Climate change is expected to have adverse effects on plant growth and development, and on pollens. Pollen grains have been widely studied in the fields of forensics, ecology, climate change and aeroallergens studies. There is an increase in allergies throughout the world especially in the industrialized countries over the past 50 years [1]. *B. papyrifera* is highly allergenic plant growing in various environmental conditions, causing frequent sneezing problems during pollination and several weeks after pollination. Variability in environmental factors induce changes in the chemical composition of lipids, carbohydrates and proteins that may increase pollen allergies [1, 2, 3]. Pollen grains are small in size 12 µm-17µm, remain in air for longer periods, and transported to longer distances [5], that make the pollen study more complex. Therefore, climate change effects on pollen grains is important to understand pollens development, and plants pollen associated allergies [6].

Climate factors that might effect pollen development requires overall assessment of lipids, carbohydrates and lipids profile that constitute pollen grain and its functions [3]. Traditional approaches used to study pollens morphology, include microscopy that identify only pollen size and shape. However, identification of specific functional changes in pollen grains due to altered lipid, protein and carbohydrate constituents and their correlation for allergy prevalence should be studied. Molecular approaches used to study proteins and lipids, are not practical due to the fact that they do not study the whole molecular structure of the pollen. These molecular methods fail to associate climatic factors effects on pollen grain and different developmental stages [5, 6]. It is important to be able to sort out the factors that are associated with climate change and are likely to influence allergenic properties of pollen.

An increase in temperature affects plant morphology, physiology, and molecular structure interferes pollen metabolism and development in some species of plants [9]. Male gametophyte development needs a large number of membrane lipids and fatty acids [10]. Heat stress during pollen development affects pollen appearance, pollen quantity as well as pollen physiology [11]. However, the heat stress effect varies in plants from species to species [11, 12]. Although pollen is of high importance for plant reproduction, the human pulmonary system is effected due to their small size and mobile in nature [14]. *B. papyrifera* (Paper mulberry) a member of Moraceae (fig family) grows in humid-tropical, subtropical, and temperate regions at an elevation of 0–1500m from sea level, and reaches a height of 12 m [15]. It is native to Japan and Taiwan and was introduced into Pakistan in early 1960s [16]. The tree is anemophilous in nature and produces large quantities of pollen grains twice a year. The pollen remains in the air for a long period of time. The pollen grains of *B. papyrifera* are known for their intense allergic potential [16,17] In Pakistan, *B. papyrifera* grows predominantly in Abbottabad, Mansehra, Kotli, Islamabad, Attock, Swabi, and Peshawar district regions, having elevation from sea levels of 1256m, 1088m, 585m, 507m, 355m, 340m, and 331m respectively. There is variation in climatic factors in these regions with differences in average temperature and precipitation. This variation affects pollen counts and pollen viability. Average temperature variation of 1 °C has been found to affect the pollen count of a various plant species (Cupressaceae, Platanus, and Quercus) from 3000–9000 pollen grains [18]. Additionally, it has been studied that change in temperature and precipitation affects biophysical and chemical properties of pollen grains [3]. *B. papyrifera* produces a large number of pollen grains that exceed 45000 grains/m^3^ in the spring season (http://www.pmd.gov.pk/rnd/rndweb/rnd_new/pr.php). A protein study of *B. papyrifera* pollen grains found 9 kD, 23 kD, and 40kD size allergens that elicit the immune response of humans to initiate allergies [16].

Heat stress causes an early bursting of the tapetum layer of pollen responsible for pollen grains nutrition and development. Tapetum rich in mitochondria [19], adds to the increased production of reactive oxygen species (ROS) under heat stress due to oxidative damage [20]. ROS activation cause regulation of pollen-associated lipid metabolites (PALMs), and they have been known to evoke immune responses [5]. Phytoprostanes present in pollens generate oxygen radicals. ROS in pollen catalyse the production of oxylipids that yield a chain reaction by producing many unstable molecules that activate gene responses for enzymes responsible for external environment changes. Some of the metabolites have been known to invoke immune responses in human beings upon exposure. These compounds present in the external layers of the pollen are more involved in pollen allergies as compared to pollen allergens that are enclosed inside the pollen and are less exposed to the human immune system. Allergenic metabolites present in pollen walls contain linoleic acid, linolenic acid, prostaglandins, phytoprostanes, sphingolipids, oxylipins, glycolipids, and their derivatives [8, 21].

The study of climate change impacts on pollen grains through reliable and user-friendly approaches is of critical importance. Fourier Transform Infrared Spectroscopy (FTIR) has been used in studying biochemical composition of plant extracts and pollens [22, 23]. The infrared waves in FTIR analysis interpret the biochemical composition of pollens. These waves by passing through the pollens, change the bonding energies of biomolecules, making the structure of pollens. Specific bond oscillations are detected by the FTIR device in the form of a spectrum. It provides an easy and quick alternative approach to study the biochemical structure of pollens [24]. Pollen sample study in its pure form through its molecular bonding energy enables FTIR to be used for studying environmental stresses effect on the pollen sample. Similarly, Liquid chromatography mass spectrometry (LCMS) determines the presence of various chemical compounds in a sample by combining chromatography and mass spectrometry. The combined use of LCMS and FTIR tools to study chemical composition of pollens in response to climate change impacts, is an ideal and state of the art approach. Therefore, the current investigations aim inter-seasonal and inter-regional variations in *B. papyrifera* pollens under varied climatic conditions.

## Materials and Methods

### Study area characteristics

The study area is comprised of four regions of Pakistan (Islamabad, Abbotabad, Kotli and Peshawar districts) having different geographical and climatic conditions. These areas are at the verge of climate shift. Islamabad is located at 507m, Abbottabad at 1256m, Kotli at 609m, and Peshawar at 331m elevation from sea level. Distance between Peshawar and Islamabad is 187 km; Peshawar and Kotli is 316 km; Peshawar and Abbottabad is 204 km; Abbottabad and Islamabad is 137 km; Abbottabad and Kotli is 265 km, and Islamabad and Kotli is 132 km. Mean annual temperature during 2010–2020 for Islamabad, Kotli and Peshawar was 21.39 °C, 22.07 °C and 23.06 °C respectively and mean annual precipitation during the period was 110.34 mm, 102.445 and 42.31mm respectively (https://www.pmd.gov.pk/en/). Abbottabad located on higher elevation that makes it cooler than all other regions and has higher precipitation than other regions. General observation supports this fact but PMD has no weather station in Abbottabad region to specify mean annual temperature and precipitation. Islamabad region has moderate environment conditions as compared to other regions of the study. It is taken as a reference to correlate inter–regional and inter–seasonal environmental effects on pollens and of other regions having variant environmental conditions.

### Pollen Sampling

Pollen of *B. papyrifera* was collected from Abbottabad and Islamabad in the summer 2020. In the spring of 2021 samples were collected from Kotli, Islamabad, and Peshawar regions during peak pollination season. Sterilized 50 ml falcon tubes were used to store flowers to avoid external sources contamination of the pollen. The samples were dried for 72 hours in shade at room temperature. Dried pollen grains were released from the samples by tipping the dry panicle of *B. papyrifera* on a 500µm pore size mesh. The isolated pollen grain was then passed through a 100µm pore size mesh to collect pure pollen grains. The pure pollen samples were stored at room temperature (28°C) in sterile 1.5ml Eppendorf tubes.

### Microscopy of pollen samples

Pollen samples were soaked in isopropanol for 10 minutes to extract lipids before applying to glass slides [25]. A small drop of suspended pollen sample was put on the glass slide and allowed dry. A drop of preheated glycerine (50 °C) was added to the pollen inserted on the spot and covered with a glass coverslip before gently heating for 15 seconds to fix the pollens.

### Fourier transform infrared analysis of pollen

20 mg of dried pollen grains were taken for each sample were placed on a FTIR IRTracer-100 knob for analysis [24]. Measurements were taken with a Vertex 70v FT-IR Spectrometer using the attenuated total reflectance (ATR) method. Infrared radiations in the range of 500-4500 cm^-1^ and a resolution of 4 cm^-1^ were selected to draw peaks [23]. The peaks obtained were normalized and data files were imported in CSV format that was processed for drawing FTIR peaks through Origin pro-2018, 32 bit.

### Liquid chromatography mass spectrometry (LCMS) analysis of pollen extracts

100 mg of dried pollen sample was taken in a 15ml falcon tube, and 2 ml of methanol was added to the tube that was kept on a shaker at room temperature for 1 hour. Then the pollen extract was passed through the SPE (Solid Phase Extraction) column through the device and pure pollen extract samples were collected in 2.0ml Eppendorf tubes. Pollen extract was subject to LCMS analysis and the compounds profile obtained were analyzed as described by Kang et.al [26].

## Results

Microscopic imaging at 100x showed that *B. papyrifera* pollen are round to oval in shape, with scar–like structures on the outer surface of the exine layer protecting the intine layer and the pollen. Summer pollen differs from spring pollen in shape (Fig 1). The pollens from different climatic zones differ in form and bio–molecular composition. Pollen FTIR analysis in its pure form showed variations in the protein region amide-II and amide–I, and lipid region. Environmental conditions of the three regions sampled varied in terms of mean annual temperature and mean annual precipitation (S1 fig, S2 fig).

**Fig 1.**
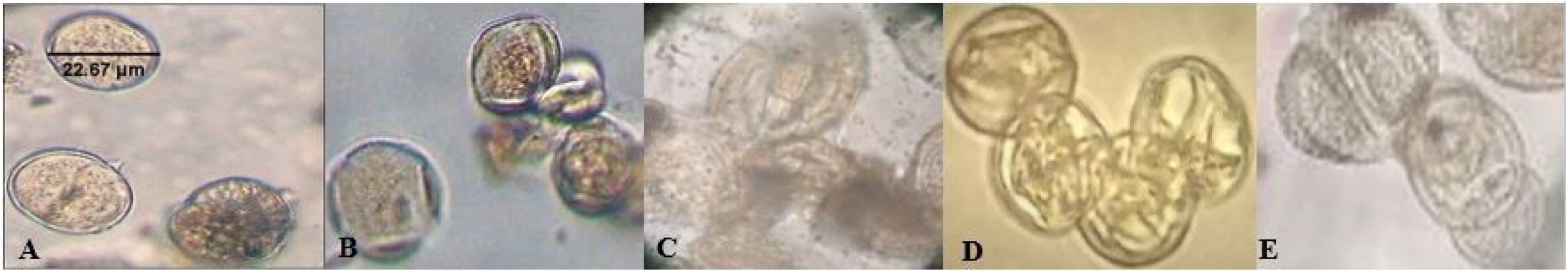
Light microscope images of *B. papyrifera* pollens captured at 100 X. A) pollen collected in Islamabad in summer 2020. B) Shows pollen collected in Abbottabad in summer 2020, C) pollens collected in Islamabad in spring 2021. D) pollen collected in Peshawar in spring 2021, and E) shows pollen collected in Kotli in spring 2021.

The LCMS analysis of summer pollens 2020 showed 33 different organic compounds. These compounds were from six organic groups where alkaloids were abundant. Terpenes, alkanes, and straight chain fatty acids (SCFA) were at the lowest concentration in the pollen samples. Four compounds were unsaturated fatty acids (succinic acid, adipic acid, pimelic acid, and sebacic acid) which have been reported to have significant role in pollen allergies [27].

The summer pollens (Fig. 1 A and B) has a rough exine shown as a dotted appearance, while spring pollens (Fig.1 C, D and E) have a smooth exine. In summer the ambient temperature is higher which seems to cause an increase in the protein quantity. Spring temperatures are lower correlating with an increase in carbohydrate and lipid content [28].

### Fourier Transform Infrared Spectroscopy (FTIR)

FTIR analysis of pollen (Fig. 2A) collected from Abbottabad and Islamabad was analysed in the summer of 2020 using spectral waive-length of 500–4000nm. Spectral regions of protein (1700–1500cm^-1^), and lipid (2900–2700cm^-1^) were analysed. The region between 1750cm^-1^ to 2750cm^-1^ showed more differences as compared to the other spectral regions. Fig. 2B shows FTIR spectra of pollen samples collected from Kotli, Islamabad, and Peshawar region in spring 2021. The spectra of the Peshawar region pollen samples had greater variation compared to spectra of pollen from Islamabad and Kotli. Also, the spectral comparison of summer and spring samples show differences in the three regions associated with chemical bonds for proteins and lipids and are listed in table 1.

**Table 1.** Spectral zone, functional group frequencies, and associated chemical bonds.

| Spectral zone | Peak frequency $\text{cm}^{-1}$ | Chemical bonds |
| --- | --- | --- |
| Lipids | 2930 | CH<sub>3</sub> stretching (lipids) |
|  | 2850 | CH<sub>2</sub> stretching (lipids) |
| Amide I<br>1700-1600$\text{cm}^{-1}$ | 1620 β-sheet<br>1650 α-helix<br>1670 β-turn | NH<sub>2</sub> bending |
| Amide II<br>1600-1500 $\text{cm}^{-1}$ | 1550 | N-O stretch |

The table shows spectral zones of lipids and proteins along with peak frequencies and chemical bonds [29].

**Fig 2.**
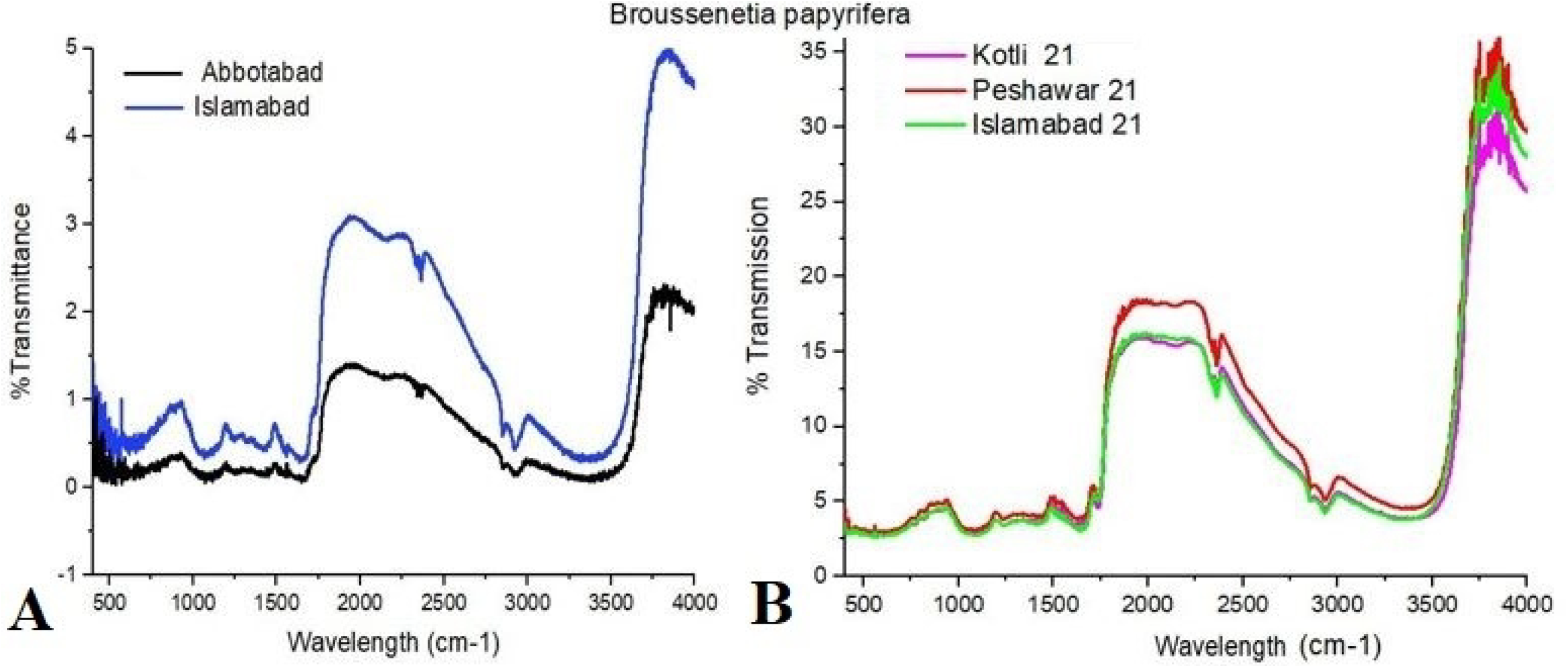
*B. Papyrifera* inter–regional and inter–seasonal pollens FTIR analysis. A) Shows pollen spectra of Islamabad and Abbottabad regions collected in summer of 2020. B) pollen spectra of Kotli, Peshawar, and Islamabad regions collected in the spring of 2021.

**Fig 2. Location effect on FTIR spectra of pollen protein content, collected in the summer of 2020 and spring of 2021**. A) Shows protein spectra of Abbottabad and Islamabad pollen collected in 2020. B) spectra of Kotli, Islamabad, and Peshawar sampled in the spring 2021.

Summer season FTIR analysis of Abbottabad and Islamabad pollens of *B. papyrifera* grown in natural environmental conditions have some variation in weather conditions reflected in the spectra of protein profile. Amide-II region of the two trees of the same species at 1540 cm^-1^ wavelength show differences in their transmittance and peaks produced. The pollen from the Islamabad region has a higher transmittance value of 1.27% at 1540 cm^-1^ while the pollen the pollen of Abbottabad has a 1.1 % transmittance at the same wavelength. Similarly, the graph peaks at the amide-II vary from one another. Amide-I region of protein at 1650 cm^-1^ wavelength has variation in the peaks of both trees pollen and their transmittance values. Pollen samples of Islamabad had higher transmittance of 1.25% as compared to pollen sample of Abbottabad 1.06% for the amide-I region.

### Pollens Principal component analysis (PCA) of different regions and different seasons

PCA of *B. papyrifera* pollens FTIR results from four different regions collected in summer and spring show variation in the five data sets. The five samples collected from four distinct regions in different seasons have variable environmental and seasonal conditions. Changes in regions and environmental factors affect pollen composition as shown in the PCA results. Only B and F datasets vectors are in the same direction that suggests similarities in the datasets. These two datasets represent Kotli and Islamabad’s spring 2021 pollens having almost similar weather conditions. Other datasets shown by vectors C–E for Peshawar March 2021 pollen sample, Islamabad August 2020 pollen sample, and Abbottabad August 2020 pollen sample, have differences. Principal Component Analysis showed that differences in climate conditions and locality affect pollen grains metabolic profile (Fig. 5).

### Light Chromatography Mass Spectrometry (LCMS) Analysis

LCMS analysis of *B. papyrifera* pollens showed different compounds which are divided into 7 groups. Among these compounds, 4 compounds (succinic acid, hexyl 2-phenylethyl ester adipic acid, di (2-phenylethyl) ester, pimelic acid, di (phenethyl) ester, and sebacic acid, 2-methylbenzyl undecyl ester) have a role in human immune system activation by eliciting interleukin-13 [21]. Other compounds are categorized as alkaloids, other lipids, carboxylic acids, alkanes, straight-chain fatty acids (SCFA), terpenes and unsaturated fatty acids (USFA). Alkaloids occur in high numbers (12) followed by other lipids (9) and carboxylic acids (5). Alkanes, SCFA, and terpenes are found very little in quantity. There was only 1 compound of straight–chain fatty acids (SCFA) (C_30_H_52_O_2_), 1 alkane (C_21_H_23_O_3_P), and 1 terpene (C_21_H_26_N_2_O_5_). Unsaturated straight-chain fatty acids (USFA) – Succinic acid, hexyl 2-phenylethyl ester (C_18_H_26_O_4_), Adipic acid, di (2-phenylethyl) ester (C_22_H_26_O_4_), Pimelic acid, di(phenethyl) ester (C_23_H_28_O_4_), and Sebacic acid, 2-methylbenzyl undecyl ester (C_29_H_48_O_4_) were also identified in the target pollens (S1Table).

## Discussion

Climate change has significant impacts on pollens biochemical composition of *B. papryrifera*. The FTIR analysis showed spectra between 4000–400 cm^-1^ that covered both fingerprint region and functional group region. Infrared light transmitted through pollens obtained pollens spectra, that analysed all compounds present in the pollen regions and showed differences in the form of peaks. These spectra for the same plant species distribution in different climatic regions showed variations in the peak height and peak area for protein Amide-II and Amide-I regions. Similarly, variations in spectra peaks were observed for the same species pollens collected in summer 2020 and spring 2021 in the same climatic region. These findings demonstrate that the pollen samples may possess quantitative differences in their biochemical composition. Pollen structure comprises of lipids, carbohydrates, and proteins [30–34]. The *B. papyrifera* pollen grains FTIR analysis, for lipids and proteins functional groups showed inter– seasonal and inter–regional differences in pollen grains (Fig 2). CH_2_ and CH_3_ stretching of lipids at 2850 cm^-1^ and 2930 cm^-1^ differ in peaks heights for pollens collected in summer 2020 in Abbottabad and Islamabad districts. The two regions have variations in temperature and precipitation that may have a possible effect on the pollen biochemical composition. Peshawar region has different environmental conditions in comparison to Islamabad and Kotli, and the respective pollens spectra varied in protein regions (1500cm^-1^–1700cm^-1^). The region comprises of Amide I region (1600 cm^-1^-1700 cm^-1^), and the Amide II region (1700 cm^-1^–1600 cm^-1^). For Amide II, peaks area at 1550 cm^-1^ for functional group [Nitroso (NO) stretch] varies. Amide I region peaks area at 1620 cm^-1^, 1650 cm^-1^, and 1670 cm^-1^ for β-sheet, α-helix, and β-turn differ from one another. Similarly, the NO-stretch peak area for spring 2021 for all three pollen samples is different. Peaks area in the Amide I region for β-sheet, α-Helix, and β-turn are similar for Kotli and Islamabad regions, and vary for the Peshawar region (Table1, Fig3). These results are consistent to previous findings which showed temperature and precipitation effects on pollen chemical composition and viability [35,36]. Period of high temperature cause bursting of tapetum layer that results in pollen sterility [37]. The current research has correlated different elevations and temperature condition effects on pollens biochemical composition through FTIR analysis, which showed significant differences in pollens collected at different elevations and different temperature conditions. Pollen tolerance to variation in temperature and elevation has been found to affect pollens viability negatively [36]. Seasonal and climatic effects on pollens are evident from pollen microscopy images (Fig 1), and FTIR spectra. Summer pollen samples spectra (Fig 2A) differ from spectra of spring pollens (Fig 2B). The protein region of Amide-I (β-sheet, α-helix and β-turn) has variation at functional group of NH_2_ bending, and Amide-II region has difference in the functional group of NO–stretch. Similar findings of heat stress effects have been studied in wheat pollen, where heat stress altered pollen lipid composition by remodelling of extraplastidic phospholipids. Heat stress decreased the quantity of more unsaturated extraplastidic phospholipids, and increased the quantity of less unsaturated extraplastidic phospholipids [2]. Moreover, variation in environmental conditions have been found to change lipids, carbohydrates and proteins expression in 813 pollen specimens from 300 distinct plant species grown in 5 different pollen seasons [38]. Similarly, the current study of pollen protein analysis through FTIR showed inter–regional and inter– seasonal differences. Amide-II peak of NO–stretch is at 1550cm^-1^ for Abbottabad and Islamabad August 2020 pollens. The same region FTIR peak in March 2021 pollens is at 1542cm^-1^. Amide-I peak for β-sheets is at 1620cm^-1^ in August 2020 pollens while the same peak for March 2021 samples is at 1618cm^-1^. α-helix peak for August 2020 pollens is at 1650cm^-1^ while the same peak for March 2021 samples is at 1648cm^-1^. β-turn for August 2020 pollens is at 1670cm^-1^ while for March 2021 pollens peaks were recorded at 1664cm-1 for all three samples. These inter–seasonal and inter–regional differences in the protein region of FTIR peaks are due to variation in temperature and precipitation conditions (S1 fig, S2 fig, Fig3). Similarly, FTIR spectroscopy has found adaptive variations in the chemical composition of proteins, lipids and carbohydrates in pollens of *Poa alpine, Anthoxanthum odoratum*, and *Festuca ovina* grown in different geographic and climatic conditions [39]. These investigations support our argument that increase in temperature and difference in elevation alter pollen chemical composition. The mean annual temperature of Islamabad region is high in comparison to average temperature of Abbottabad, and the mean annual precipitation of Islamabad is low in comparison to average precipitation of Abbottabad. Similarly, spring 2021 pollen analysis showed variation in protein regions in the FTIR spectra. Peshawar region has extreme climatic conditions (high temperature and less precipitation) in comparison to Islamabad. Islamabad and Kotli regions have moderate variations in weather conditions and elevations. The spectra analysed for the three regions show elevated peaks and enhanced transmission for Peshawar as compared to Kotli and Islamabad. This enhancement in the spectra for the Peshawar region can be correlated to the weather conditions of Peshawar (S1 fig). The high–temperature stress role in in-vitro pollen germination and pollen tube growth in *Pisum sativum L*. modification has been verified [40]. The FTIR peaks in the proteins and lipids functional groups region collected during different seasons and from different regions varied in terms of infrared light transmission and peak area (Fig 3, Fig 4).

**Fig 3.**
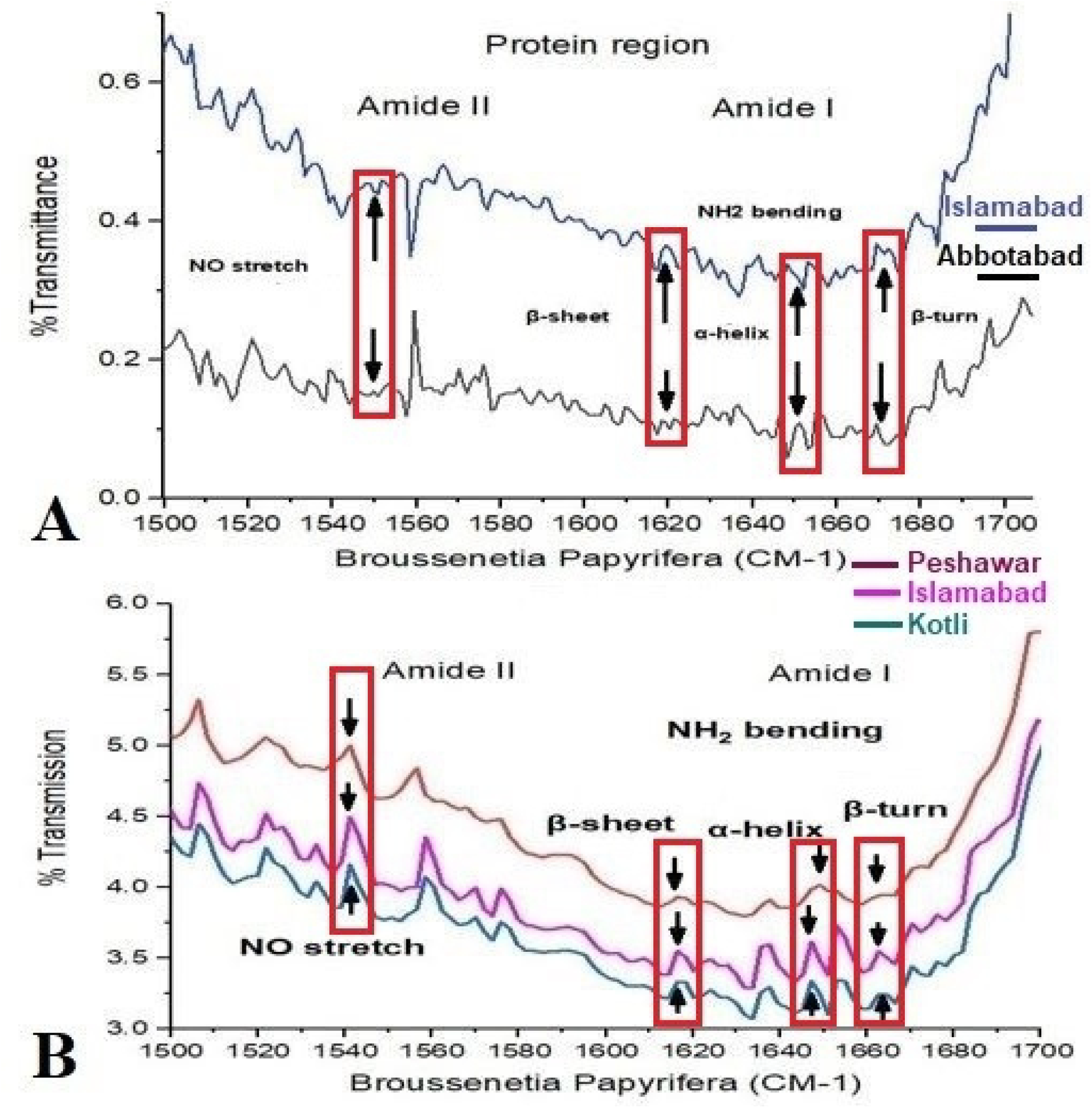
Seasonal and climatic effects on lipid region of pollens. A) Shows summer 2020 pollen FTIR spectra for lipid region 2840cm^-1^ to 2940cm^-1^. B) Show spring 2021 pollen FTIR spectra for lipid region 2840cm^-1^ to 2940cm^-1^.

**Fig 4.**
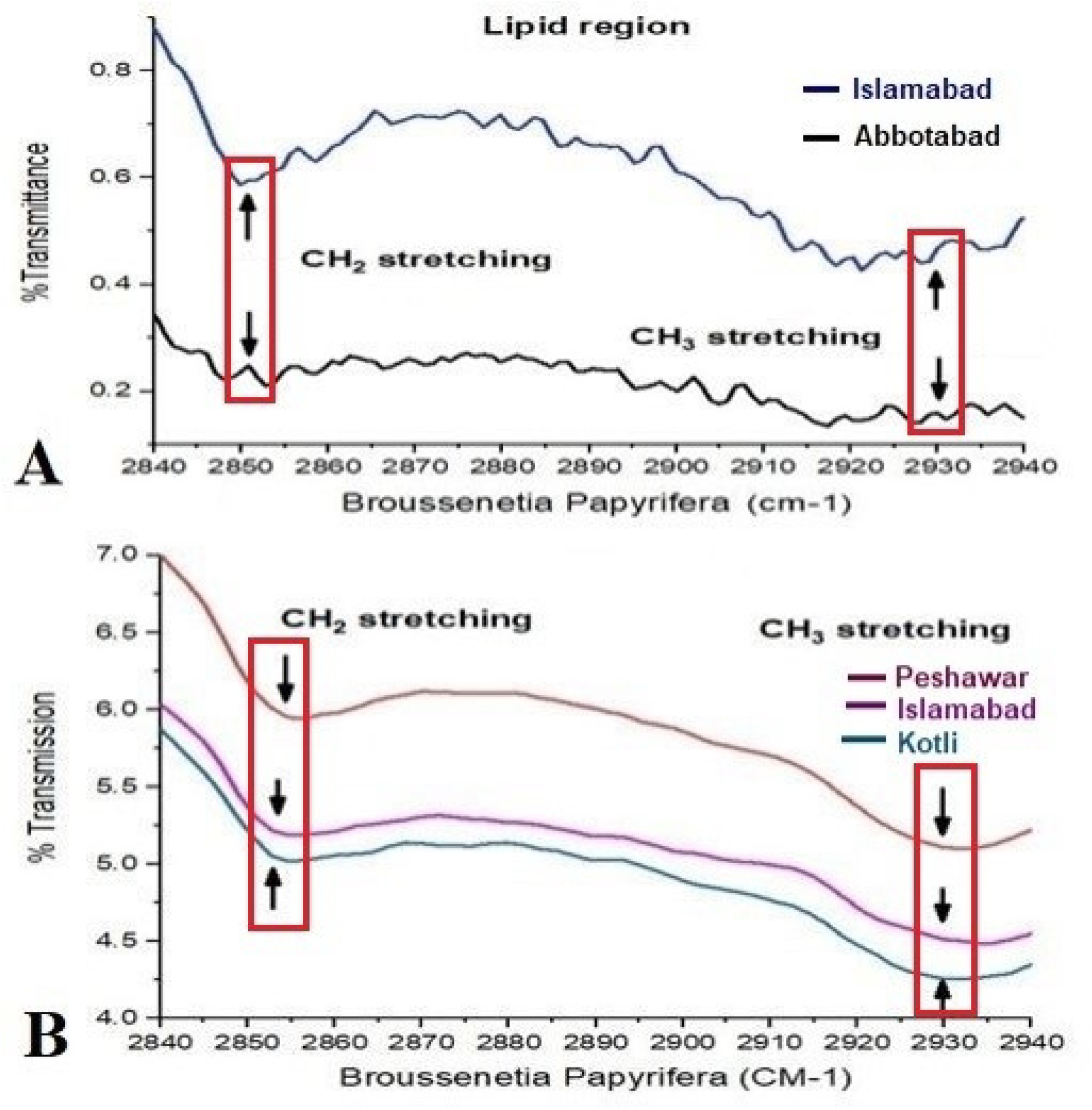
Principal component analysis (PCA) of *B. papyrifera* pollens from four different regions. *B. papyrifera* FTIR transmittance of pollen. A *–* E. show pollen from Kotli 2021, Peshawar 2021, Islamabad 2020, Abbottabad 2020, and Islamabad 2021 respectively. PCA analysis shows differences in the pollen FTIR datasets of different regions and different seasons.

**Fig 5.**
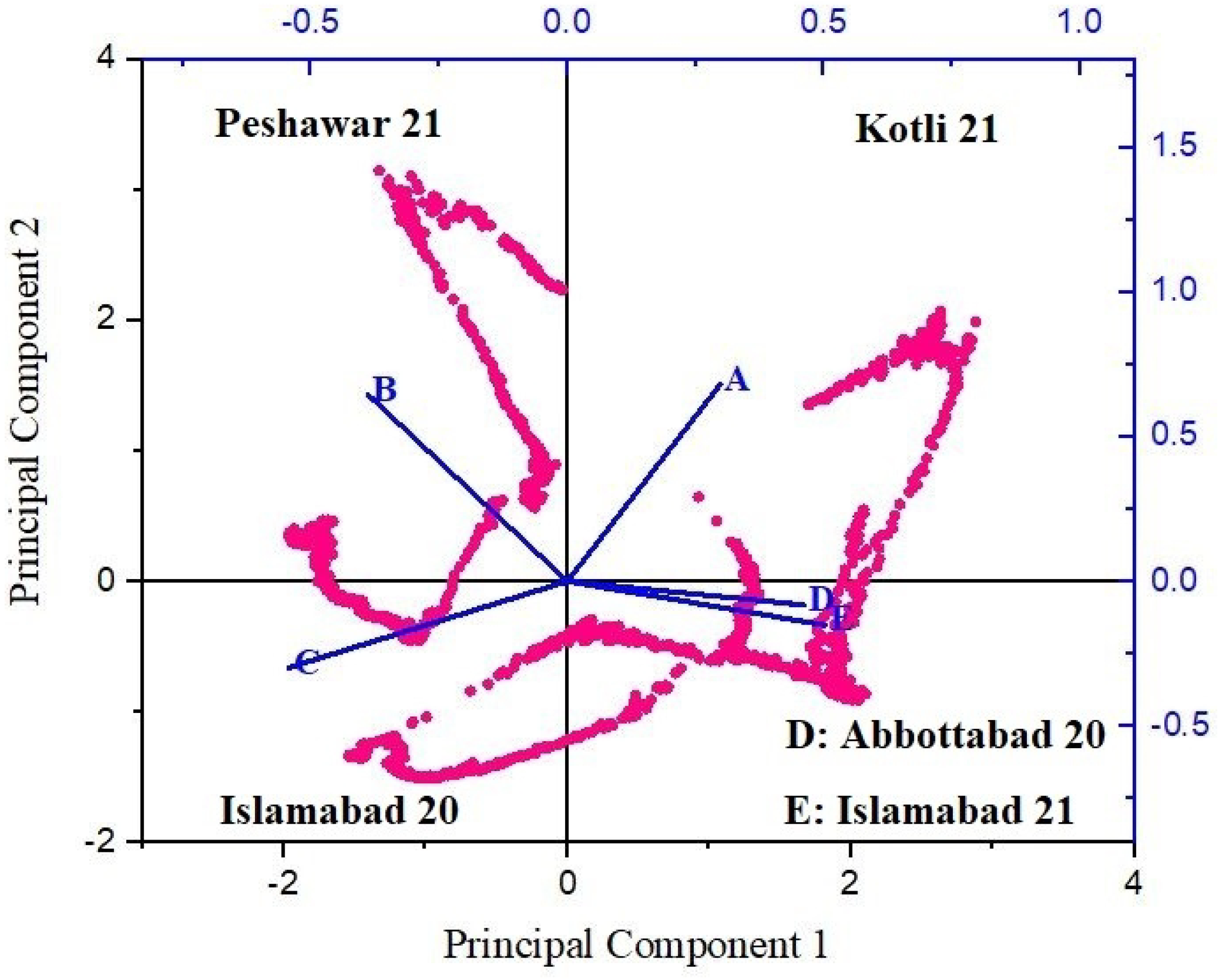
LCMS analysis of *B. papyrifera* pollens. Y-axis shows charge to mass ratio for the ions of various compounds separated at different time intervals while X-axis denotes acquisition time in minutes. The sample run time is 17 minutes having the highest peak at 13^th^ minute 12.9369. The analysis shows variance in charge to mass ratios of ions present in the pollens that make unique peaks. Peaks analysis show presence of different compounds in the analysed pollens (for details see S1Table)

**Fig 6.**
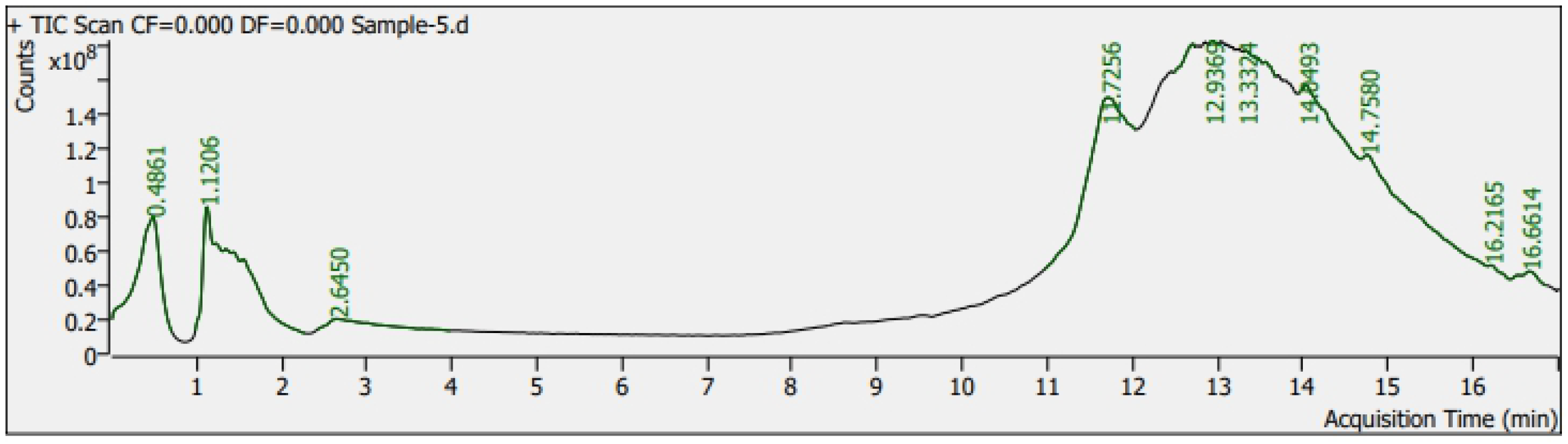
Schematic distribution of organic compounds in *B. papyrifera* pollen grains by LCMS analysis. Total compounds identified in the pollen samples are classified into 7 groups: alkanes, alkaloids, carboxylic acids, terpenes, other lipids, straight chain fatty acids, and unsaturated fatty acids (USFA). USFA marked in red color shows the presence of allergenic compounds in the pollen grains of *B. papyrifera*. The pollen contains 4 unsaturated fatty acids that induce T-helper-2 (TH2) cells associated Interleukine-13 (IL-13) which are involved in nasal allergies in human [21].

The difference in temperature, precipitation and geographic location correlated with differences the spectral region of *B. papyrifera*. As variation in temperature and precipitation act as stress that activate an oxidative stress response in plants that alter pollen metabolism and development. High–temperature stress to pollens yields an increase in pollen proteins while low–temperature stress causes enhancement of pollen lipids and carbohydrates [39,41]. This phenomenon is reflected in the pollen spectra of *B. papyrifera* collected in spring 2020 from Islamabad and Abbottabad having variance in environmental conditions (average temperature and average precipitation) (Fig 2). The temperature difference in the two regions can have a possible effect on the FTIR spectra of the pollen samples. Increase in temperature and drought has been found to affect pollen weight, protein content and allergen contents [42]. In our studies, inter–regional and inter–seasonal pollens studied through Principal Component Analysis (PCA) showed significant differences in the FTIR datasets of pollens, which also confirmed differences in pollens chemical composition. In a recent study, honeybee pollens PCA analysis has shown significant differences in pollens biochemical composition [43], which further strengthen findings of our research that pollens chemical composition varies with change in elevation and temperature.

The *B. papyrifera* produces allergenic pollens that cause hypersensitivity in human beings as these pollens are small in size (12-17 µm) and are capable to pass through human nasal cavity, and reach deep into the lungs. Two allergens in *B. papyrifera* pollens have been identified [16]. Our investigations on *B. papyrifera* pollens metabolome analysis through LCMS have found presence of succinic acid, hexyl 2-phenylethyl ester (C_18_H_26_O_4_), adipic acid, di (2-phenylethyl) ester (C_22_H_26_O_4_), pimelic acid, di (phenethyl) ester (C_23_H_28_O_4_), and sebacic acid, 2-methylbenzyl undecyl ester (C_29_H_48_O_4_). These compounds have significant role in causing immune system elicitation and allergies [5]. Exposure of epithelial layers of the deep nasal cavity to such small pollens that possess both allergens and allergenic metabolites may be involved in the incitation of the human immune system. We assume that this can be the possible reason for the high pollen allergies incidences in Islamabad and surrounding regions during the flowering season where daily pollen count in the air exceeds 45000 grains/m^3^ [44].

The morphological studies and biochemical analysis of pollens through FTIR show that environmental variables (temperature and precipitation), have significant effect on pollen proteins and lipids contents. Such climate roles affecting pollens biochemical composition have been verified in several studies [33,45–48], and increasing prevalence of pollen allergenicity [49]. Findings of this research are unique which may help in determining the temperature effects on activation of reactive oxygen species (ROS), heat shock proteins (HSP), and oxidative stress mechanisms in pollens. The temperature effect has been studied to affect the viability of pollen grains [50]. Pollen samples analysis through FTIR present a new approach to study temperature effect on the overall development and metabolism of pollen grains [39]. Pollen–associated lipid metabolites (PALMs) have potential roles in pollen metabolism and development, and have been found to elicit TH2 cells [51] and IL13 that cause allergies [21]. In our findings, the FTIR analysis showed variation in the lipid region peaks for lipids specific functional groups (CH_3_, CH_2_), lipids region peak height and peak area (Fig 4). Moreover, the *B. papyrifera* pollens analysis through LCMS has identified 33 compounds from 7 different groups (Fig 7). Out of these 7 groups, USFA group has 4 compounds whose role has been established in allergies [21,52]. Variation in climatic factors that may act as stresses, activates PALMs that incite the human immune system.

**Figure7.**
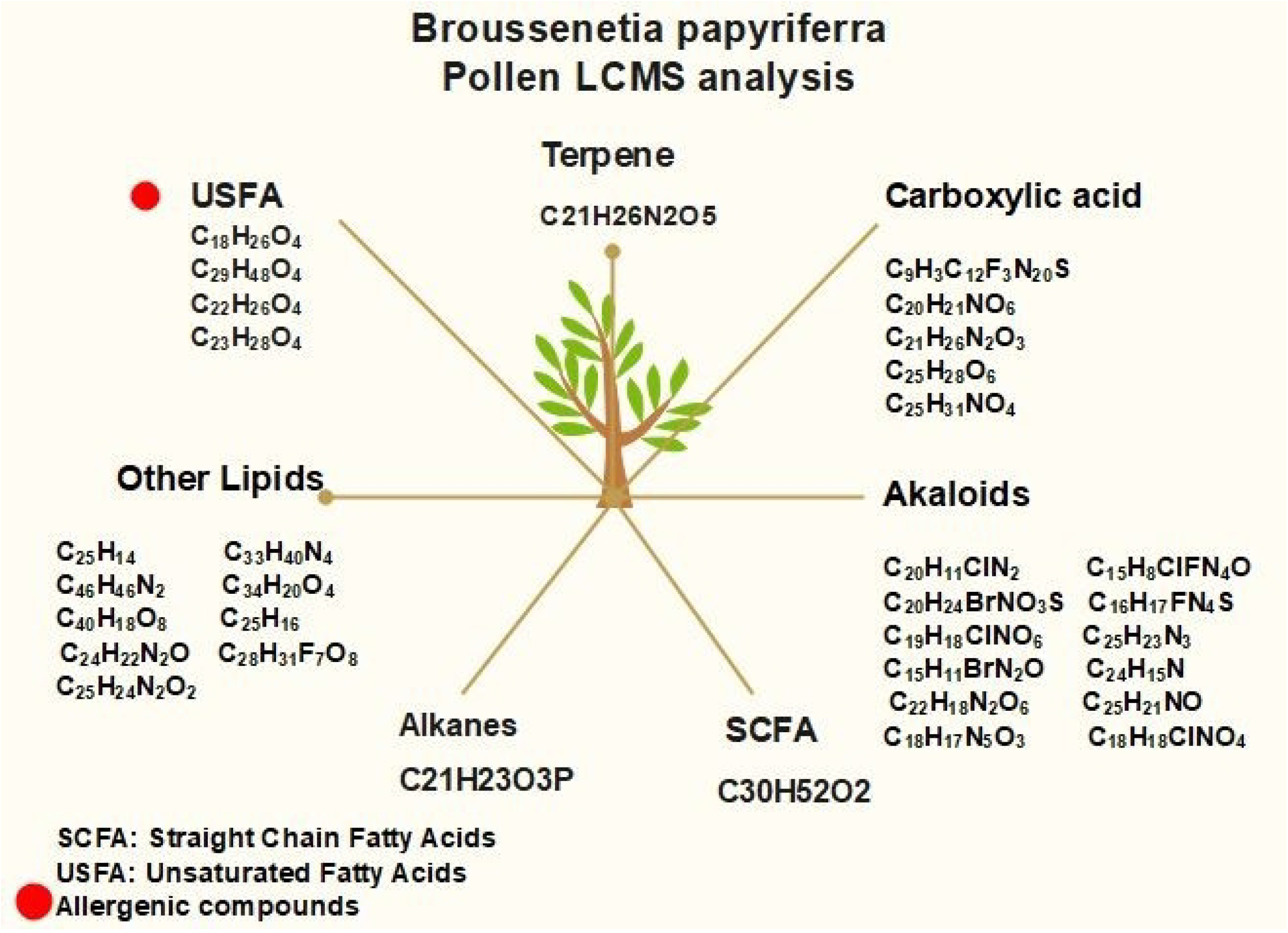

## Conclusion

Climate change can have significant impacts on pollen morphology, pollen viability, and biochemical structure of *B. papyrifera*. Like other plants, the pollens of *B. papyrifera* are also subject to detrimental effects of climate change. An increase in temperature causes premature bursting of pollen tubes and activation of ROS that damage pollens by oxidative damage. This study has found that increase in temperature, decrease in precipitation, and difference in elevations yield variation in *B. papyrifera* pollen grains appearance and biochemical structure. This difference in biochemical structure occurs both in variation in climate variables within the same season and in different seasons. Similarly, climate factors affect pollens morphology and molecular composition based on locality. *B. papyrifera* pollen grains contain 33 different organic compounds. Among these organic compounds, 4 unsaturated fatty acid compounds have a possible role in the elicitation of the human immune system in the form of allergies. The findings are unique which may provide a baseline model system for predicting variation in pollen structure and associated allergies for other species in response to climate shift.

## Acknowledgements

We are thankful to Higher Education Commission (HEC) Pakistan for providing financial support to conduct this study under the umbrella of HEC-NRPU Project No. 8231 titled “Strategic development of base line protocols to predict climate change impacts on plants and prevalence of associated allergies in Pakistan”. We are also thankful to Climate Data Processing Centre Pakistan Meteorological Department, Pakistan for providing precipitation and temperature data.

## Authors Contribution

MH performed the experiment and wrote the manuscript, SN helped in data compilation and analysis, and wrote the manuscript. RG helped in experimental design, reviewed the manuscript and improved the English language of the manuscript. ZA conceptualized the idea, designed the experimental strategies, reviewed the manuscript and supervised the whole project.

## Supporting Information

**S1 Fig. Mean annual temperature of Kotli, Peshawar and Islamabad from 2010 to 2020**. The bar graph shows mean annual temperature of three regions. Highest temperature for Peshawar and Kotli regions was 23.8 °C and 22.6 °C in the year 2016, and for Islamabad it was 22.5 °C in 2010. The lowest mean annual temperature for Islamabad was 20.4 °C in 2019, for Kotli it was 21.5 °C in 2014 and for Peshawar region, it was 22.6 °C in 2019.

**S2 Fig. Mean annual precipitation of Kotli, Islamabad and Peshawar from 2010-2020**. Peshawar region has highest mean annual precipitation in 2014 and lowest in 2015 while Kotli region has highest mean annual precipitation in 2015 and lowest in 2018. Similarly, Islamabad region has highest mean annual precipitation in 2013 and lowest in 2017.

